# Gonadotropin-releasing hormone increases the release of extracellular vesicles from gonadotropes

**DOI:** 10.1101/2020.09.19.304592

**Authors:** Numfa Fungbun, Makoto Sugiyama, Ryota Terashima, Shiro Kurusu, Mitsumori Kawaminami

**Author notes:** Corresponding author Faculty of Veterinary Medicine, Okayama University of Science, Ikoinoka 1-3, Imabari, Ehime 794-0885, Japan. The authors declare no competing financial interests.

## Abstract

While extracellular vesicles play a role in intercellular communication, it is not known how their release is regulated. We show here that hypothalamic gonadotropin-releasing hormone (GnRH) stimulates extracellular vesicle (EV) formation associated with annexin A5 (ANXA5) from pituitary gonadotropes. The results show that 1) membrane blebs containing ANXA5 are produced after GnRH agonist (GnRHa) stimulation of gonadotropes and that this is observed *in situ* as a loss of distinction at cell-cell boundaries, 2) EV containing ANXA5 are increased by GnRHa, 3) a gonadotrope-derived EV fraction stimulates LH release from other gonadotropes, and 4) an increase in ANXA5-loaded EV occurs in the plasma of ovariectomized rats. Finally, we also showed that 5) GnRHa stimulation of blebbing and EV-ANXA5 were suppressed by a protein kinase A inhibitor. These present results demonstrate a novel autocrine/paracrine mechanism mediated through the production of EV containing ANXA5. A hormonal regulation mechanism of cell-cell communication by means of EV is suggested.

---

Annexin A5 (ANXA5) is a member of the annexin family, which consists of twelve calcium-dependent phospholipid binding proteins (ANXA1-A11 and A13) (Pepinsky *et al*., 1988; Liemann and Lewit-Bentley, 1995; Moss and Morgan, 2004). All annexins share a similar structure of four repeats of approximately 70 amino acids and a highly conserved sequence, with the exception of eight repeats in ANXA6 (Pepinsky *et al*., 1988; Liemann and Lewit-Bentley, 1995; Moss and Morgan, 2004). We previously found that hypothalamic GnRH stimulates the expression of ANXA5 in the pituitary gonadotrope (Kawaminami *et al*., 2002a; Kawaminami *et al*., 2002b; Kawaminami *et al*., 2008). Furthermore, we found that ANXA5 augments gonadotropin secretion and enhances the stimulatory effect of GnRH (Kawaminami *et al*., 2002a). However, it is still unclear how ANXA5 augments gonadotropin release.

The primary sequences of annexins do not show signal sequences (Pepinsky *et al*., 1988; Moss and Morgan, 2004). Although they have been observed in the cytoplasm, as expected, the annexins also have been found extracellularly (Pepinsky *et al*., 1988; Van Eerden *et al*., 2006; Ravassa *et al*., 2007). As an example of an extracellular action of ANXA5, we found abundant ANXA5 on the endothelium in the placenta and demonstrated that ANXA5 on the wall of vessels protects mothers from unexpected blood coagulation and pregnancy loss (Ueki *et al*., 2012). Another family member, ANXA1, is externalized by folliculostellate cells by means of the ABC transporter (Wein *et al*., 2004). However, it is unknown whether and how ANXA5 is externalized by gonadotropes.

Cell-to-cell communication via hormones/cytokines is a prerequisite for synchronized functions in multicellular organisms. Various mechanisms for communication have been found in mammals, including endocrine, juxtacrine, autocrine, and paracrine. The secretion of bioactive substances and the expression of cognate receptors on target cells are necessary for this kind of communication. Hence, hormones, neurotransmitters, cytokines and growth factors would be categorized all in a same group based on this point. Intercellular communication occurs also by gap junctions between cells, as seen in cardiomyocytes and smooth muscle cells. Recently, novel communication mechanisms via extracellular vesicles (EV), exosomes and ectosomes or microvesicles, are attracting attention (Valadi *et al*., 2007; Cocucci and Meldolesi, 2015). An exosome is released by fusion of a multivesicular endosome and has a particle size of less than 100 nm (Raposo and Stoorvogel, 2013; Cocucci and Meldolesi, 2015). Usually, an exosome fraction is included in small extracellular vesicles (S-EV) obtained by ultracentrifugation at more than 100,000 xg (Dujardin *et al*., 2014). On the other hand, an ectosome or microvesicle is produced by pinching off a vesicle from a region of the extruded membrane (Raposo and Stoorvogel, 2013; Cocucci and Meldolesi, 2015). The size of these structures is more than 100 nm, and they can be found in large extracellular vesicles (L-EV) obtained by centrifugation at 20,000 xg (Dujardin *et al*., 2014). We report here that L-EV containing ANXA5 are produced by the blebbing of GnRH-stimulated gonadotropes and that such L-EV augment luteinizing hormone (LH) secretion. This is the first report, as far as we know, that reveals the hormonal control of L-EV formation, through which the regulation of the surrounding cells is attained.

## Materials and Methods

### Animals

Adult female Wistar Imamichi rats were maintained in light (5:00-19:00 h) and temperature (23±3°C) controlled rooms. They were fed laboratory chow and tap water *ad libitum*. All experimental procedures and animal care were performed according to the guidelines for animal treatment of Kitasato University and approved by the Institutional Animal Care Committee at Kitasato University (15-032).

### Cell culture

#### Primary culture of pituitary cells

Primary cultures of anterior pituitary cells were prepared from adult female Wistar Imamichi rats as reported previously (Kawaminami *et al*., 2002a; Kawaminami *et al*., 2002b). Anterior pituitary glands were cut into 1 mm^3^ pieces and dispersed with 0.25% trypsin (Invitrogen, NY) and 10 mM EDTA (Dulbecco’s modified Eagle medium; DMEM, pH 7.4) for 40 minutes at 37°C with slow stirring by means of a spinner flask. Pituitary tissue pieces were rinsed and mechanically disrupted by passage through flame-polished Pasteur pipettes. The cells were washed and then resuspended at 10^6^ cells/ml in DMEM supplemented with 10% fetal calf serum (Gibco Life Technologies, NY), antibiotic-antimycotic mixture (Gibco Life Technologies) and nonessential amino acids (Gibco Life Technologies). The cells were maintained in an atmosphere of 95% air, 5% CO_2_ and 100% humidity at 37°C.

#### LßT2 cells

A gonadotrope-derived cell line, LßT2, was a kind gift of Dr. P Mellon (UC San Diego). LßT2 was cultured in Dulbecco’s modified Eagle medium with high glucose (Invitrogen, Tokyo, Japan) supplemented with 10% fetal calf serum and 1% antibiotic-antimycotic mixture (Gibco Life Technologies) (Kawaminami *et al*., 2008). The cells were grown in 75 cm^2^ flasks and maintained in an atmosphere of 95% air, 5% CO_2_ and 100% humidity at 37°C. The cells were subcultured prior to reaching 80% confluency.

### Hormone assay

LH levels in the medium were measured by means of a time-resolved fluorometric immunoassay with rat LH assay kits (National Hormone and Peptide Program, NHPP) and the Delfia system (Perkin Elmer, Waltham, MA) as reported previously (Kawaminami *et al*., 2002a). LH-I-9 was labeled with europium using a Delfia Eu-labeling kit (PerkinElmer). Anti-rabbit gamma globulin goat serum was prepared in our laboratory and purified by ammonium sulfate precipitation. A 96 well immunoplate (Nunc, Tokyo, Japan) was coated with anti-rabbit gamma globulin, after which an anti-rat LH serum (1:4,000 dilution) was layered over. The sample medium was incubated, followed by the addition of optimally diluted Eu-labeled hormone, which were then incubated at 4°C overnight. The specific fluorescence was measured with an ARVO multi label reader (Perkin Elmer) after adding enhancer (Perkin Elmer).

### Immunocytochemical staining

Cells grown on poly-L-lysine-coated cover slips were fixed with 4% PFA for 15 minutes and rinsed with cold acetone. The fixed cells were incubated with 3% fetal calf serum for 1 hour at room temperature. The immunocytochemical staining of LßT2 cells with a rabbit anti-rat ANXA5 antibody (AB_2827744, 1:10,000) was performed and detected by an Alexa 488 goat anti-rabbit IgG. Negative controls without a primary antibody were prepared, and the absence of nonspecific fluorescence was confirmed (data not shown). Actin was stained with phalloidin (Thermo Fisher Diagnostics) to make the cell shape clearer. Double immunocytochemical staining of primary cultures of pituitary cells was performed using a rabbit anti-rat ANXA5 antibody (1:10,000) and a guinea pig anti-rat LHβ antibody (1:10,000, NIDDK-anti-betaLH-IC-2). The secondary antibodies used were Alexa 488 goat anti-rabbit IgG and Alexa 568 goat anti-guinea pig IgG (Thermo Fisher Diagnostics, Tokyo, Japan). The lack of cross-reactivity of the antibodies was also confirmed (data not shown). The nucleus was stained with DAPI (VECTASHIELD Mounting Medium with DAPI, Funakoshi, Tokyo, Japan). The immunocytochemical staining of the ANXA5 distribution was observed by a confocal laser microscope (Zeiss 710, Jena, Germany).

### Transmission electron microscopic (TEM) observation of pituitary tissues

Hemi-pituitary glands were incubated with or without the GnRH agonist (GnRHa, 100 nM) for 10 and 30 minutes. The tissue samples were fixed by immersion in Karnovsky solution (2% glutaraldehyde, 2% paraformaldehyde in 0.05 M cacodylate buffer, pH 7.4) (Sugiyama *et al*., 2019). The specimens were trimmed to a size of approximately 1 mm^3^ and immersed in the same Karnovsky solution. Tissue samples were postfixed with 1% osmium tetroxide/1.5% potassium ferrocyanide. The tissue samples were then dehydrated and embedded in epoxy resin. Ultrathin sections were cut using an Ultracut N (Reichert-Nissei, Wein, Austria), stained using uranyl acetate followed by lead citrate and examined using a Hitachi H-7650 transmission electron microscope (Hitachi Ltd, Tokyo, Japan). The particulate fractions were fixed and stained with phosphotungstic acid on a carbon-coated copper grid and observed by using TEM (80 MeV, × 15000) as reported previously (Otani *et al*., 2019).

### Isolation of membrane particles by sequential centrifugation

The medium was collected and placed immediately on ice. After separating the cells and cellular debris by centrifugation at 300 xg, 4°C for 10 minutes (multipurpose refrigerated centrifuge LX-120; Tomy, Tokyo, Japan), the supernatant was then centrifuged at 2,000 xg, 4°C for 20 minutes (high speed refrigerated centrifuge Suprema 21; Tomy, Japan) to pellet apoptotic bodies. Subsequently, the supernatant was centrifuged at 20,000 xg, 4°C for 45 minutes (high speed refrigerated centrifuge Suprema 21) to collect the membrane particles, namely, L-EVs. The supernatant was then ultracentrifuged at 110,000 xg, 4°C for 60 minutes to isolate S-EV (Optima XL-80K, Beckman Coulter Lifesciences, Tokyo, Japan). The plasma samples were centrifuged similarly but without the 300xg centrifugation.

### Western blotting

Proteins were separated by SDS-PAGE using 12% poly-acrylamide gel. Electrophoretic transfer to a PVDF membrane was performed. The membrane was blocked with 5% skim milk and 3% fetal calf serum for 1 hour at room temperature. Immunodetection of ANXA5 was accomplished with an anti-ANXA5 antibody (1:10,000) and peroxidase-labeled anti-rabbit IgG (1:50,000). Chemiluminescence was detected by an ECL Prime detection system (GE Healthcare, Tokyo, Japan) and an ImageQuant LAS 4000 (GE Healthcare). Each band was analyzed and shown as a fold increase in ANXA5 intensity. We did not employ any internal standard protein for Western blotting analysis but standardized samples by number of cells and volume. It is because there was no appropriate standard protein for EV and we thought the number of EV itself would change.

### Experimental designs

#### Immunocytochemical observation of ANXA5 in cells stimulated by GnRHa

LßT2 cells were seeded on poly-L-lysine-coated cover slips at 100,000 cells per cover slip. The next day, the medium was replaced with or without 100 nM of GnRHa for 10 and 30 minutes. The GnRHa was fertirelin acetate (Concereal^®^ injection) purchased from Nagase Medicals (Itami, Japan). The cells were fixed and processed for immunocytochemical staining using an anti-ANXA5 antibody. Primary cultures of anterior pituitary cells were prepared. After cell dissociation, the pituitary cells were seeded at 100,000 cells on poly-L-lysine-coated cover slips. Two days after cell seeding, the medium was changed to medium with or without 100 nM of GnRHa, followed by incubation for 10 or 30 minutes. Pituitary cells were fixed and subjected to double immunocytochemical staining with anti-ANXA5 and anti-LHß antibodies.

#### Detection of ANXA5 in the particulate fraction of the medium of cultured LßT2 cells

LßT2 cells were maintained as described above, seeded on 10 cm culture-dishes. The experiment was performed after the cells grew to 80% confluency. The medium was removed and cells were gently rinsed with prewarmed serum-free medium. LßT2 cells (4 dishes each) were incubated with or without 100 nM GnRHa and maintained for 10, 30 or 180 minutes. Each medium was collected and placed immediately on ice. The particulate fractions were separated by sequential centrifugation of the medium. Pellets from the 20,000 xg and 110,000 xg centrifugations were suspended in 25 µl of sample buffer. The suspension was subjected to SDS-PAGE and Western blotting with an anti-ANXA5 antibody.

### The effect of L-EV on LH release

LßT2 cells were grown and maintained as described. The L-EV fractions were isolated from the conditioned medium of LßT2 cells, after incubation with 100 nM of GnRHa for 3 hours, by centrifugation at 20,000 xg. The L-EV fraction of the control incubation was also prepared. A portion of the control pellet was further preincubated with GnRHa for 3 hours (pellet incubated with GnRHa). Preincubation without GnRHa was also performed. Each conditioned pellet was washed with serum-free medium and then centrifuged at 20,000xg for 45 minutes before examining the effect on LH release in cultured LßT2 cell (96 well format) for 24 hours of incubation. The LH levels in the medium samples were measured by time-resolved fluorometric immunoassay.

### Electron microscopic observation of hemi-pituitary tissue stimulated by GnRHa

The anterior pituitary glands of adult female Imamichi rats were utilized in the study. The anterior pituitary glands were cut at the isthmus to make two equivalent halves. The hemi-pituitary glands were incubated with or without 100 nM GnRHa for 10 and 30 minutes. Pituitary tissues were subjected to observation with a transmission electron microscope as described above.

### Detection of ANXA5 in the plasma particulate fraction of ovariectomized rats

Adult female Wistar Imamichi rats were used. They were sham operated (n=3) or ovariectomized (OXV, n=3), and after seven days, the blood was collected from the aorta to heparinized syringe under isoflurane anesthesia. After centrifugation of the blood at 2,000 xg, 4 ml of plasma was centrifuged sequentially at 20,000 xg and 110,000 xg. Each pellet fraction was washed with cold PBS and centrifuged again at 20,000 xg for 45 minutes and 110,000 xg for 60 minutes, respectively. The washed pellets isolated from the plasma of sham or ovariectomized rats were resuspended with sample buffer. Twenty microliters of each suspension was subjected to SDS-PAGE and Western blotting with an anti-ANXA5 antibody. For analyzing the plasma ANXA5 content, 200 μl of 110,000 xg supernatant was resuspended in 4 times the volume of cold acetone for 2 days at −20°C. The precipitate was collected by centrifugation at 2,500 rpm. The protein pellets were suspended with sample buffer, and 20 μg protein/20 μl sample buffer was subjected to SDS-PAGE and Western blotting.

### Effects of inhibitors

LβT2 cells were challenged with various inhibitors of signal transduction. A GnRH antagonist (cetrorelix, 100 nM), PKC inhibitor (GF 109203x, 10 nM), MAPKK inhibitor (PD 98059, 28 μM) or PKA inhibitor (H89, 30 μM) were added and incubated for one hour. The culture medium was then changed to a medium containing each inhibitor with or without GnRHa (100 nM), followed by incubation for 30 minutes.

### Statistical analysis

Each value is presented as the mean±SEM. Statistical analysis was performed using the Student’s *t* test for the comparison of two groups and the Tukey test for multiple comparisons. P values less than 0.05 were considered to be significant.

## Results

GnRHa induced obvious membrane blebbing that contained ANXA5 on the surface of cells of the gonadotrope cell line, LßT2, within 10 minutes (Fig. 1A-b). ANXA5-positive blebs were also observed after 30 minutes of GnRHa administration (Fig. 1A-c).

**Fig. 1.**
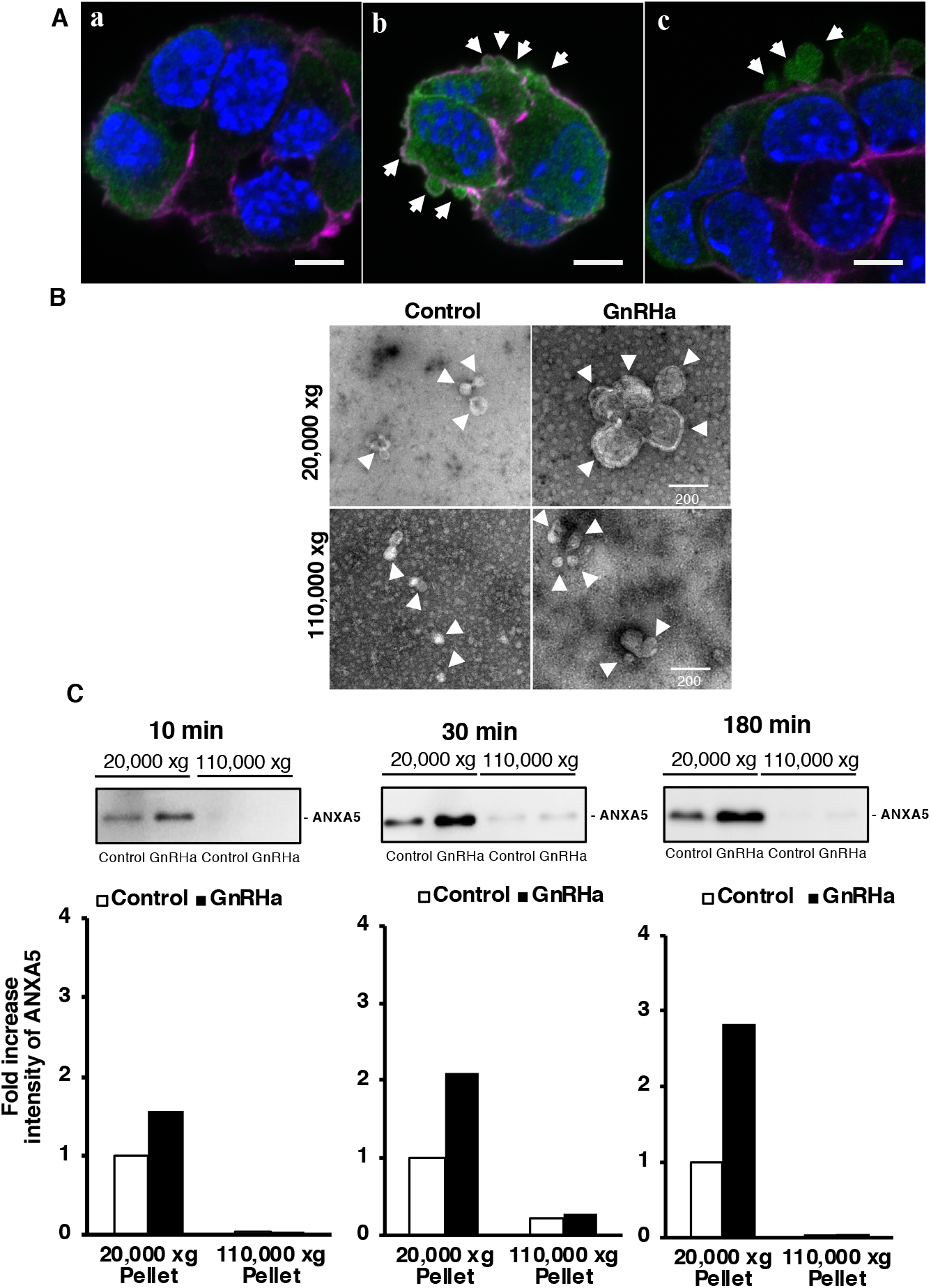
GnRH agonist induces blebbing containing ANXA5 in LβT2 cells A: Immunocytochemical staining for ANXA5 in LßT2 cells. The cells were subjected to immunocytochemistry after GnRH agonist (100 nM) treatment. The arrows indicate the ANXA5-containing blebs. Green: ANXA5; magenta: actin; blue: DAPI. (a) Control incubation and (b) 10 and (c) 30 minute incubations with the GnRH agonist. The scale bar is 5 µm. The experiments were repeated three times. B: The conditioned medium of LßT2 cells treated with GnRH agonist (100 nM for 30 minutes) was fractionated by sequential centrifugation. The membrane particles obtained by 20,000 xg and 110,000 xg centrifugation were observed with a transmission electron microscope after negative staining. White arrow heads indicate the particles. The scale bar is 200 nm. C: Western blotting for ANXA5 in the 20,000 xg and 110,000 xg pellets was performed. The 20,000 xg and 110,000 xg pellets were isolated from the conditioned medium of LßT2 cells after treatment with or without the GnRH agonist for 10, 30 and 180 minutes. Each band was analyzed and is shown as a fold increase. ANXA5 was detected obviously in the 20,000 xg pellet and increased gradually after 10, 30 and 180 minutes of GnRH agonist treatment. This experiment was repeated twice.

Membrane particles in the conditioned medium of LßT2 cells were isolated by sequential centrifugation. Pellets obtained by centrifugation at 20,000 xg and 110,000 xg were negatively stained and observed by transmission electron microscopy (Fig. 1 B). GnRHa increased the number of particles in the 20,000 xg pellet, and the diameter of those particles ranged from 150 to 200 nm, matching the reported size of microvesicles (Raposo and Stoorvogel, 2013; Dujardin *et al*., 2014). Smaller particles of less than 100 nm were seen in the 110,000 xg pellet, consistent with their identification as exosomes; in this case, there was no obvious difference between the control and the GnRHa-treated sample.

Particulate fractions were subjected to Western blot analysis with an anti-ANXA5 antibody. Conditioned medium from LßT2 cells was collected after 10, 30 and 180 minutes of GnRHa stimulation. Particulate fractions were separated by sequential centrifugation at 2,000, 20,000 and 110,000 xg. Fractions pelleted at 20,000 and 110,000 xg were analyzed (Fig. 1 C). ANXA5 was observed predominantly in the 20,000 xg pellet, and the amount seen increased after only 10 minutes of GnRHa treatment. GnRHa continued to increase the amount of ANXA5 in the 20,000 xg pellet until 180 minutes (Fig. 1C). Only scarce amounts of ANXA5 were seen in the 110,000 xg pellet, and there was no difference between the control and the GnRHa-treated samples.

Primary cultures of pituitary cells were subjected to double immunocytochemical staining with antibodies against the specific ß subunit of LH (anti-LHß (magenta) and ANXA5 (green)) (Fig. 2A). Blebs appeared on primary gonadotropes after 10 and 30 minutes of GnRHa stimulation (Fig. 2A-b and c). These blebs contained ANXA5 (Fig. 2A-b and c, arrows).

**Fig. 2.**
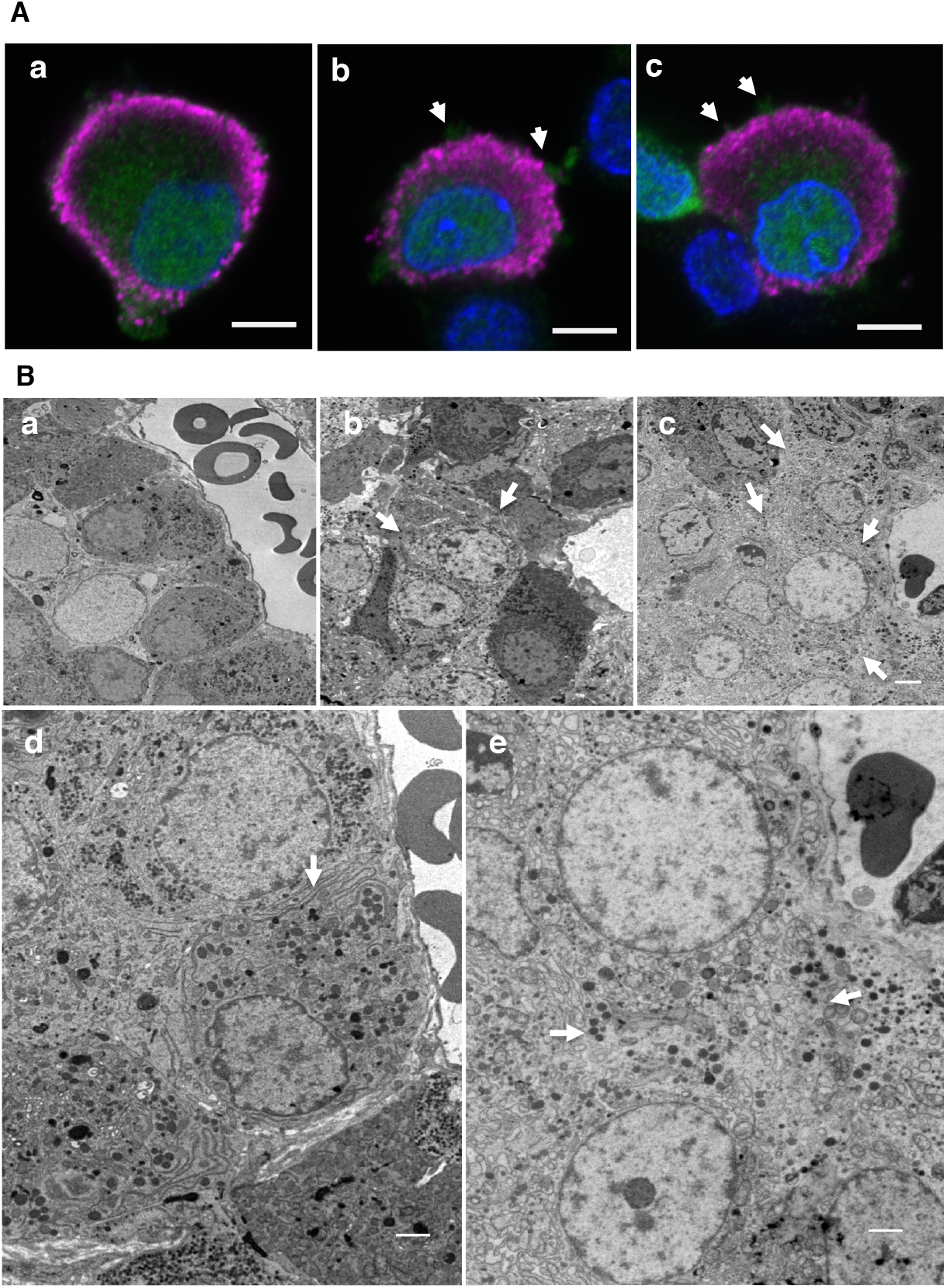
GnRH agonist stimulation of ANXA5-containing blebbing in primary gonadotropes A: A primary culture of pituitary cells were subjected to double immunocytochemical staining with anti-LHβ and anti-ANXA5 antibodies after GnRH agonist (100 nM) administration. The arrows indicate the ANXA5-containing buds on the gonadotropes. Green: ANXA5; magenta: LHβ; blue: DAPI. The scale bar is 5 µm. B: Hemi-pituitary glands were incubated with or without 100 nM GnRHa for 10 and 30 minutes. (a) Control incubation and (b) 10 and (c) 30 minute incubations with the GnRH agonist; (d) and (e) are higher magnifications of the control incubation and the 30 minutes incubation with the GnRH agonist, respectively. Note that the boundary of the cells became obscure after GnRH agonist administration and that many particles appeared (Fig. 2-b, c, e arrows compared to Fig. 2-d (arrows)). The scale bar is 2 µm. This experiment was repeated twice.

Hemi-pituitary organ culture was performed, and the pituitary glands were fixed after 30 minutes of GnRHa stimulation. The pituitary sections of the GnRHa-stimulated hemi-pituitaries showed obscure cell borders for morphologically defined gonadotropes (Fig. 2B, in which the arrows indicate the assumed border areas). The control pituitary tissue that was the contralateral half showed clear cell borders (Fig. 2B-d, arrow).

A L-EV fraction collected from the conditioned medium of the GnRHa-treated (100 nM, 3 hours) LβT2 cells was shown to stimulate LH release of other LβT2 cells (Fig. 3A). To determine the potential effect of carry-over of GnRHa from the producers to the responders, the control L-EVs were incubated with GnRHa for 3 hours and then used to test the responders. Although the L-EVs that absorbed the GnRHa significantly stimulated LH release, the extent was clearly smaller than that seen with the L-EVs from the GnRHa-treated cells.

**Fig. 3.**
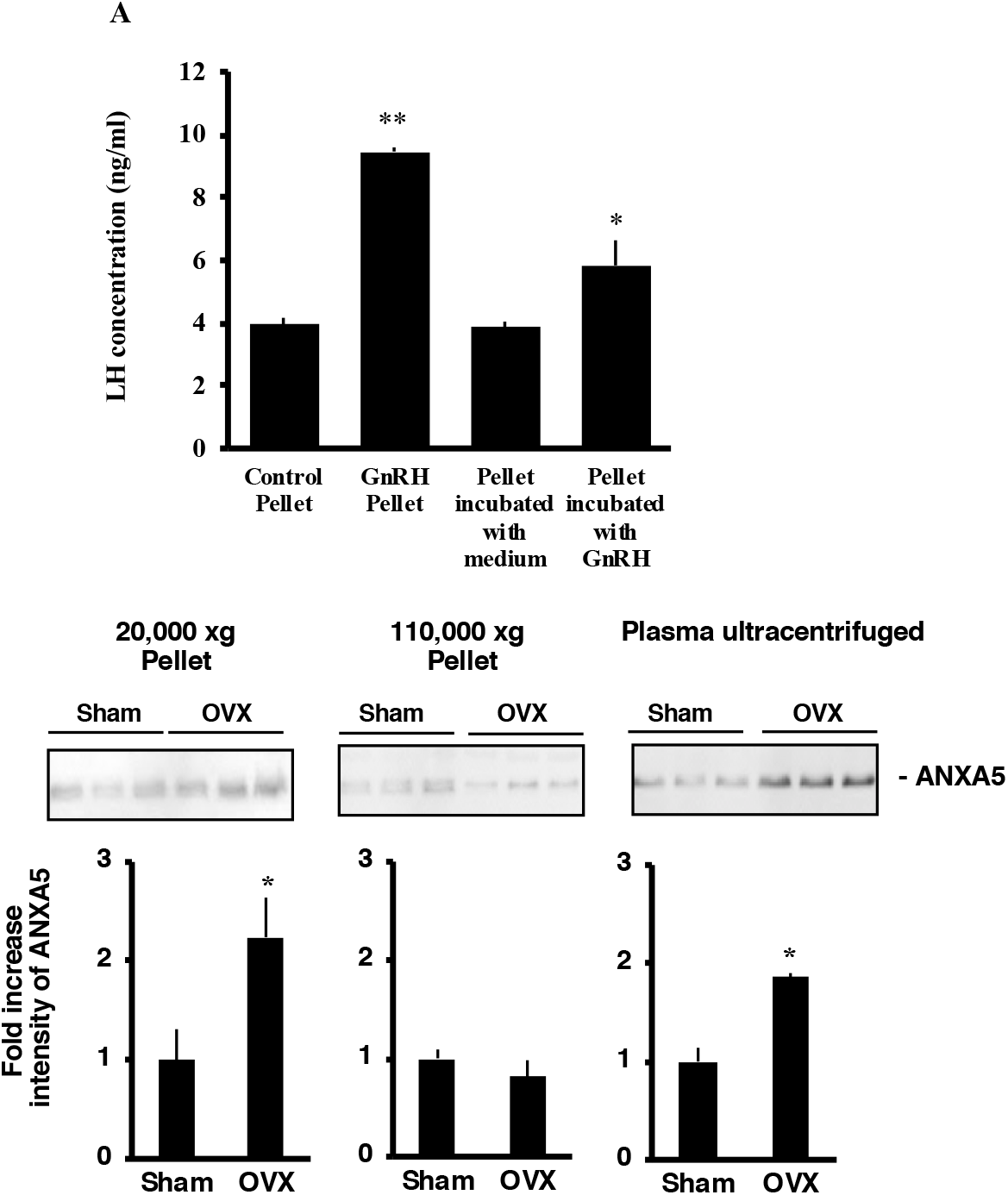
The physiology of L-EV containing ANXA5 A: The effect of L-EV on LH release. The L-EV fraction prepared from the conditioned medium of LßT2 cells after GnRH agonist treatment was added to the culture of LßT2 cells and the effect on LH release was examined. The conditioned medium of LßT2 cells treated with 100 nM GnRH agonist for 3 hours was centrifuged at 20,000 xg (GnRH Pellet). The L-EV fraction prepared from the control incubation was also prepared (control pellet). A portion of the control pellet was further preincubated with the GnRH agonist (100 nM) before examining the effect on LH release (pellet incubated with the GnRH agonist). Preincubation without the GnRH agonist was also performed (pellet incubated with medium). LH levels in the medium are presented as the mean values±SEM of five samples. The asterisks indicate a significant difference from all groups: *p<0.05, **p<0.0001. This experiment was repeated twice. B: Augmentation of L-EV containing ANXA5 in the plasma of ovariectomized rats. The 20,000 and 110,000 xg pellets were isolated from the plasma of sham or ovariectomized (OVX) rats (n=3) after 7 days. The pellet was subjected to Western blotting for ANXA5. Densitometric analysis was performed, and the intensity of each band is shown as the mean±SEM of the fold increase. The asterisk indicates a significant difference from each sham group: * p<0.05.

ANXA5-containing L-EVs were increased in the plasma of ovariectomized rats (Fig. 3B). Blood plasma was obtained from female rats seven days after ovariectomy. L-EVs and soluble fractions were prepared by sequential centrifugation and subjected to SDS-PAGE and Western blotting for ANXA5. Ovariectomy significantly augmented the ANXA5 content in the 20,000 xg pellet (Fig. 3B). Free ANXA5 was also increased in the soluble fraction that was obtained as the 110,000 xg supernatant from the ovariectomized rat plasma (Fig. 3B). ANXA5 in the 110,000 xg pellet was very low and was not changed by ovariectomy.

As the signal transduction of the GnRH receptor is reported to be coupled with protein kinase C, MAPK and adenylate cyclase (Perrett and McArdle, 2013), the effects of these inhibitors on bleb formation and ANXA5 content in L-EVs were examined. The GnRH antagonist cetrorelix clearly inhibited the effect of GnRHa on bleb formation and ANXA5 accumulation in the L-EVs fraction (Fig. 4A-c yellow arrow, B-c). Inhibition of PKC and MAPKK showed no obvious effect on bleb formation or the amounts of ANXA5 by Western blot (Fig. 4A-d, e, B-d, e). H89 led to the disappearances of ANXA5 in the blebs (Fig. 4A-f, yellow arrow). H89 reduced ANXA5 in the L-EV fraction (Fig. 4B-f).

**Fig. 4.**
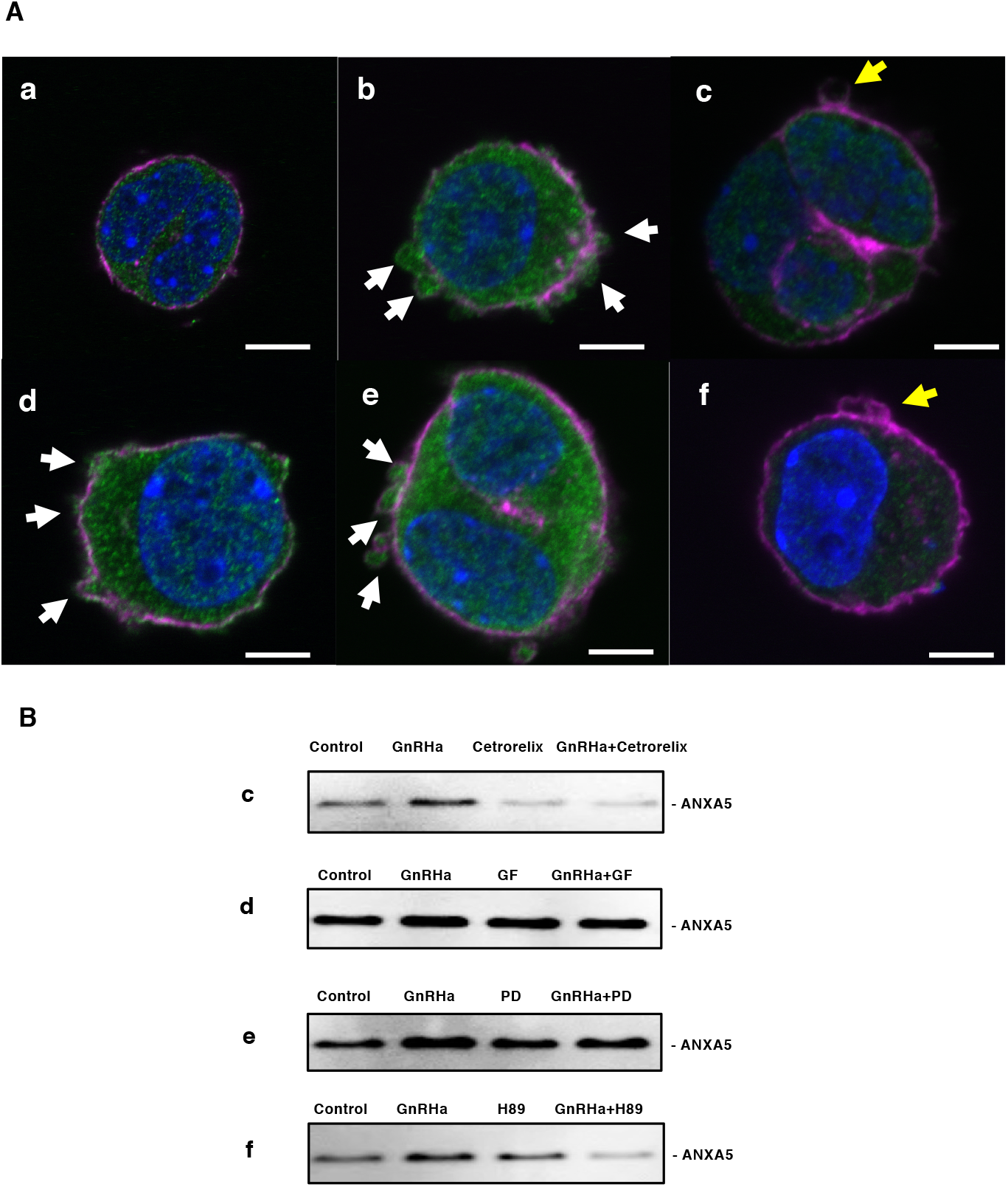
The effect of signal transduction inhibitors on bleb formation and ANXA5 excretion (A) Immunocytochemical staining for ANXA5 in LßT2 cells treated with GnRH agonist and inhibitors. a) Control incubation, b) GnRH agonist, c) cetrorelix (GnRH antagonist), d) GF (protein kinase C inhibitor GF109203x), e) PD (MAPKK inhibitor PD98059), and f) H89 (protein kinase A inhibitor H89). A one hour incubation with inhibitors was followed by 30 minutes of incubation with GnRHa. Green: ANXA5; magenta: actin; blue: DAPI. (B) Western blotting for ANXA5 in the L-EV fraction The L-EV fraction was obtained from the conditioned medium by centrifugation (20,000 xg) and was subjected to Western blot analysis of ANXA5. c: Cetrorelix, GnRH receptor antagonist; d: GF, protein kinase C inhibitor GF109203x; e: PD, MAPKK inhibitor PD98059; and f: H89, protein kinase A inhibitor H89. The series of experiments was repeated twice.

## Discussion

In the present study, we determined that abundant blebs appeared on the surface of gonadotropes shortly after GnRHa administration. This phenomenon has already been reported by others, and in that report, the blebs were presumed to be retracted and thought to contribute to cell movement (Navi *et al*., 2017). In the present study, bleb formation was found to accompany the formation of L-EVs. Evidence in support of this statement includes the appearance of ANXA5-containing blebs following GnRHa stimulation, as shown by immunocytochemical staining and the concurrent production of ANXA5-containing L-EVs in the 20,000 xg fraction, as shown by electron microscopy with the negative staining of vesicles and Western blotting of the L-EV fraction. Thus, although it is not known whether the blebbing and L-EV formation are directly related phenomenon, both are induced by GnRH stimulation. It is suggested that ANXA5 is released from cells by this L-EV formation mechanism. This phenomenon is similar to apocrine secretion seen in some exocrine glands.

Initial experiments used the gonadotrope cell line, LßT2. To be sure that these cells were representative of normal pituitary gonadotropes in regard to bleb formation, primary cultures were examined, in which gonadotropes were identified by immunocytochemical staining with an anti-LHß subunit antibody. These cells also produced blebs in response to GnRHa stimulation. To be sure that blebbing was not a feature of cells in a two-dimensional culture, electron microscopic examination of GnRHa-treated hemi-pituitaries was performed. In this instance, the gonadotropes were detected by their morphology, as described in classical texts (Kurosumi *et al*., 1991). Dramatic changes in the border of putative gonadotropes were seen after GnRHa stimulation, suggesting that *in situ* gonadotropes form blebs and L-EVs in response to GnRHa.

Since ovariectomy removes negative feedback from ovarian hormones on the hypothalamic pituitary axis, the GnRH level in portal blood is presumed to be very high at seven days after ovariectomy (Wheaton and McCann, 1976). Thus, the production of ANXA5-containing L-EV was expected to be substantial, and the increase in ANXA5 in the 20,000 xg fraction would be detected peripherally in these animals. Consistent with expectations, we saw a doubling of vesicle-contained ANXA5 and an almost doubling of ANXA5 outside of vesicles in the blood plasma. At the moment, it is unclear how this extravesicular ANXA5 became extravesicular. ANXA5 release from vesicles would take place naturally, but we cannot rule out vesicular damage during centrifugation or circulation. We would say that the formation and the release of L-EVs would occur physiologically.

Extracellular vesicles (EVs) are known as two different types, namely, microvesicles (ectosomes) and exosomes. They are different in their origin, size, protein markers and method of formation (Cocucci and Meldolesi, 2015). Intercellular transport of materials by EV is reported for proteins, lipids, and RNA (Valadi *et al*., 2007; Mathivanan *et al*., 2012). ANXA5 has been reported to be present in exosomes and has been proposed to be one of the markers for exosome identification in addition to CD9, CD63, CD81, Alix, TGS101, FLOT1, FLOT2 and ANXA2 (Principe *et al*., 2013). The presence of exosomes containing ANXA5 has been suggested to be a biomarker of cancer, such as colorectal carcinoma (Valcz *et al*., 2016). We did not see much ANXA5 in the S-EV fraction in the present study. It is possible that the presence of ANXA5 in exosomes is a cancer-specific feature or may reflect a better separation of the two fractions in the present study.

Here, we have demonstrated that the excretion of ANXA5 by L-EVs, probably mainly microvesicles, constitutes a novel enhancing mechanism of responses to GnRH, but how ANXA5, carried in vesicles or in the extracellular fluid, actually increases secretion is not known. We speculate that there is a mechanism allowing ANXA5 from the vesicle to interact with the plasma membrane of the gonadotrope, where it acts as a calcium channel (Rojas *et al*., 1990; Berendes *et al*., 1993). If this is the case, extracellular ANXA5 would increase intracellular calcium and subsequently hormone release. Further study is needed to determine whether this is the case.

An increase in plasma L-EVs containing ANXA5 would be anticipated in postmenopausal women because of the reduced negative feedback to the hypothalamic-pituitary axis for gonadotropin secretion. How increased circulating ANXA5 may contribute to postmenopausal pathologies is an area ripe for further investigation.

The formation of blebs in response to GnRHa and blockade using the antagonist raises the question of what signaling pathway mediates bleb and L-EV formation after activation of the GnRH receptor. Of the inhibitors examined, only the protein kinase A (PKA) inhibitor, H89, suppressed bleb formation and the externalization of ANXA5. Suppression of bleb formation by both the GnRH antagonist and the PKA inhibitor was accompanied by reduced cytosolic ANXA5. Therefore, it is hypothesized that increased ANXA5 expression and bleb formation are controlled by the same signaling pathway initiating from the GnRH receptor. The major signal transduction pathway from the GnRH receptor is through Gq/11 (Perrett and McArdle, 2013). However, Gs has been reported to be coupled with the GnRH receptor as well (Perrett and McArdle, 2013). GnRH activates Gs-adenylate cyclase and PKA (Liu *et al*., 2002). As we have shown, activated PKA is necessary to form blebs and L-EVs containing ANXA5. It has been previously reported that the process by which GnRH induces bleb formation is through extracellular signal regulated kinase (ERK1/2) and RhoA-ROCK (Navi *et al*., 2017). We did not see an obvious tendency to reduce L-EV ANXA5 by PD98059 in the present study. As the inhibition of MEK was reported not to be sufficient to suppress bleb formation by GnRH (Navi *et al*., 2017), it appears that protein kinase A is a principal pathway used by GnRH to stimulate the formation of blebs and L-EVs.

In summary, ANXA5 of gonadotropes is externalized, primarily by L-EV formation following GnRH-cAMP signaling. L-EVs, containing ANXA5 and derived from gonadotropes, stimulate LH release from the same and/or other gonadotropes, both *in vitro* and *in vivo*. Hormonal control of L-EV formation would be a novel means of intercellular communication.

## Acknowledgements

The authors thank Dr. Ameae Walker (University of California, Riverside) for critical discussion and reading the manuscript and Ms. Miyoko Nakata for her technical support.

## Author Contributions

Numfa Fungbun conducted most parts of the experiments. Makoto Sugiyama worked with Numfa Fungbun to take the electron micrographs. Ryota Terashima and Shiro Kurusu always participated in discussions with Numfa Fungbun to accomplish this work and supported the laboratory work. Mitsumori Kawaminami is the PI and completed the manuscript.

## Author Information

Affiliation

Research Group for Animal Health Technology, Faculty of Veterinary Medicine, Khon Kaen University, Khon Kaen, Thailand

Numfa Fungbun

Veterinary Anatomy, Kitasato University, Aomori, Japan

Makoto Sugiyama

Veterinary Physiology, Kitasato University, Aomori, Japan

Ryota Terashima, Shiro Kurusu and Mitsumori Kawaminami

Present affiliation of MK

Veterinary Physiology, Okayama University of Science, Ehime, Japan

Correspondence and requests for materials should be addressed to MK.

## References

Berendes, R., A. Burger, D. Voges, P. Demange, and R. Huber. 1993. Calcium influx through annexin V ion channels into large unilamellar vesicles measured with fura-2. FEBS Lett. 317:131–143. https://doi.org/10.1016/0014-5793(93)81507-V

Cocucci, E., and J. Meldolesi. 2015. Ectosomes and exosomes: Shedding the confusion between extracellular vesicles. Trends Cell Biol. 25:364–372. https://doi.org/10.1016/j.tcb.2015.01.004

Dujardin, S., S. Bégard, R. Caillierez, C. Lachaud, L. Delattre, S. Carrier, A. Loyens, M.C. Galas, L. Bousset, R. Melki, G. Aurégan, P. Hantraye, E. Brouillet, L. Buée, and M. Colin. 2014. Ectosomes: A new mechanism for non-exosomal secretion of Tau protein. PLoS One. 9. https://doi.org/10.1371/journal.pone.0100760

Kawaminami, M., S. Etoh, H. Miyaoka, M. Sakai, M. Nishida, S. Kurusu, and I. Hashimoto. 2002a. Annexin 5 messenger ribonucleic acid expression in pituitary gonadotropes is induced by gonadotropin-releasing hormone (GnRH) and modulates GnRH stimulation of gonadotropin release. Neuroendocrinology. 75:2–11. https://doi.org/10.1159/000048216

Kawaminami, M., Y. Tsuchiyama, S. Saito, M. Katayama, S. Kurusu, and I. Hashimoto. 2002b. Gonadotropin-releasing hormone stimulates annexin 5 messenger ribonucleic acid expression in the anterior pituitary cells. Biochem. Biophys. Res. Commun. 219:915–920. https://doi.org/10.1006/bbrc.2002.6573

Kawaminami, M., N. Uematsu, K. Funahashi, R. Kokubun, and S. Kurusu. 2008. Gonadotropin releasing hormone (GnRH) enhances annexin A5 mRNA expression through mitogen activated protein kinase (MAPK) in LbetaT2 pituitary gonadotrope cells. Endocr. J. 55:1005–1014.

Kurosumi, K., H. Ozawa, K. Akiyama, and T. Senshu. 1991. Immunoelectron Microscopic Studies of Gonadotrophs in the Male and Female Rat Anterior Pituitaries, with Special Reference to their Changes with Aging. Arch. Histol. Cytol. 54:559–571. https://doi.org/10.1679/aohc.54.559

Liemann, S., and A. Lewit-Bentley. 1995. Annexins: a novel family of calcium- and membrane-binding proteins in search of a function. Structure. 3:233–237. https://doi.org/10.1016/S0969-2126(01)00152-6

Liu, F., I. Usui, L.G. Evans, D.A. Austin, P.L. Mellon, J.M. Olefsky, and N.J.G. Webster. 2002. Involvement of both Gq/11 and Gs proteins in gonadotropin-releasing hormone receptor-mediated signaling in LβT2 cells. J. Biol. Chem. 277:32099–32108. https://doi.org/10.1074/jbc.M203639200

Mathivanan, S., C.J. Fahner, G.E. Reid, and R.J. Simpson. 2012. ExoCarta 2012: Database of exosomal proteins, RNA and lipids. Nucleic Acids Res. 40:D1241–D1244. https://doi.org/10.1093/nar/gkr828

Moss, S.E., and R.O. Morgan. 2004. The annexins. Genome Biol. 5. https://doi.org/10.1186/gb-2004-5-4-219

Navi, L.R. Ben, A. Tsukerman, A. Feldman, P. Melamed, M. Tomic, S.S. Stojilkovic, U. Boehm, R. Seger, and Z. Naor. 2017. GnRH induces ERK-dependent bleb formation in gonadotrope cells, involving recruitment of members of a GnRH receptor-associated signalosome to the blebs. Front. Endocrinol. (Lausanne). 8. https://doi.org/10.3389/fendo.2017.00133

Otani, K., Y. Fujioka, M. Okada, and H. Yamawaki. 2019. Optimal isolation method of small extracellular vesicles from rat plasma. Int. J. Mol. Sci. 20. https://doi.org/10.3390/ijms20194780

Pepinsky, R.B., R. Tizard, R.J. Mattaliano, L.K. Sinclair, G.T. Miller, J.L. Browning, E.P. Chow, C. Burne, K.S. Huang, D. Pratt, L. Wachter, C. Hession, A.Z. Frey, and B.P. Wallner. 1988. Five distinct calcium and phospholipid binding proteins share homology with lipocortin I. J. Biol. Chem. 263:10799–10811.

Perrett, R.M., and C.A. McArdle. 2013. Molecular mechanisms of gonadotropin-releasing hormone signaling: Integrating cyclic nucleotides into the network. Front. Endocrinol. (Lausanne). 4. https://doi.org/10.3389/fendo.2013.00180

Principe, S., E.E. Jones, Y. Kim, A. Sinha, J.O. Nyalwidhe, J. Brooks, O.J. Semmes, D.A. Troyer, R.S. Lance, T. Kislinger, and R.R. Drake. 2013. In-depth proteomic analyses of exosomes isolated from expressed prostatic secretions in urine. Proteomics. 13:1667–1671. https://doi.org/10.1002/pmic.201200561

Raposo, G., and W. Stoorvogel. 2013. Extracellular vesicles: Exosomes, microvesicles, and friends. J. Cell Biol. 200:373–383. https://doi.org/10.1083/jcb.201211138

Ravassa, S., A. González, B. López, J. Beaumont, R. Querejeta, M. Larman, and J. Díez. 2007. Upregulation of myocardial Annexin A5 in hypertensive heart disease: Association with systolic dysfunction. Eur. Heart J. 28:2785–2791. https://doi.org/10.1093/eurheartj/ehm370

Rojas, E., H.B. Pollard, H.T. Haigler, C. Parra, and A.L. Burns. 1990. Calcium-activated endonexin II forms calcium channels across acidic phospholipid bilayer membranes. J. Biol. Chem. 265:21207–21215.

Sugiyama, M., D. Shindo, N. Kanada, T. Ohzeki, K. Yoshioka, M. Funaba, and O. Hashimoto. 2019. Inducible brown/beige adipocytes in retro-orbital adipose tissues. Exp. Eye Res. 184:8–14. https://doi.org/https://doi.org/10.1016/j.exer.2019.03.021

Ueki, H., T. Mizushina, T. Laoharatchatathanin, R. Terashima, Y. Nishimura, D. Rieanrakwong, T. Yonezawa, S. Kurusu, Y. Hasegawa, B. Brachvogel, E. Pöschl, and M. Kawaminami. 2012. Loss of maternal annexin A5 increases the likelihood of placental platelet thrombosis and foetal loss. Sci. Rep. 2. https://doi.org/10.1038/srep00827

Valadi, H., K. Ekström, A. Bossios, M. Sjöstrand, J.J. Lee, and J.O. Lötvall. 2007. Exosome-mediated transfer of mRNAs and microRNAs is a novel mechanism of genetic exchange between cells. Nat. Cell Biol. 9:654–695. https://doi.org/10.1038/ncb1596

Valcz, G., O. Galamb, T. Krenïcs, S. Spisïk, A. Kalmïr, I.V. Patai, B. Wichmann, K. Dede, Z. Tulassay, and B. Molnïlr. 2016. Exosomes in colorectal carcinoma formation: ALIX under the magnifying glass. Mod. Pathol. 29:928–983. https://doi.org/10.1038/modpathol.2016.72

Van Eerden, P., X.X. Wu, C. Chazotte, and J.H. Rand. 2006. Annexin A5 levels in midtrimester amniotic fluid: Association with intrauterine growth restriction. Am. J. Obstet. Gynecol. 194:1371–13376. https://doi.org/10.1016/j.ajog.2005.11.005

Wein, S., M. Fauroux, J. Laffitte, P. De Nadaï, C. Guaïni, F. Pons, and C. Coméra. 2004. Mediation of annexin 1 secretion by a probenecid-sensitive ABC-transporter in rat inflamed mucosa. Biochem. Pharmacol. 67:1195–1202. https://doi.org/10.1016/j.bcp.2003.11.015

Wheaton, J.E., and S.M. McCann. 1976. Luteinizing hormone-releasing hormone in peripheral plasma and hypothalamus of normal and ovariectomized rats. Neuroendocrinology. 20:296–310. https://doi.org/10.1159/000122496

